# How Electric Field Remodels the Nanofibril Structure of Chitosan Hydrogels: the Role of Dewetting During Electro-Assembly

**DOI:** 10.1101/2025.04.25.650622

**Authors:** Aarion Romany, Gregory F. Payne, Jana Shen

## Abstract

Electrofabrication has emerged as a versatile technique for creating complex functional materials from self-assembling biopolymers such as chitosan and collagen; however, a molecular-level understanding of electric cueing remains lacking. Here we investigate how a mild electric field (similar in magnitude to that imposed on biological membranes) remodels the nanofibril structure of chitosan hydrogels using all-atom molecular dynamics simulations. The simulations revealed a mechanism of active dewetting, in which the electric field enhances fibrillar order and induces compaction along the sheet-stacking direction through expulsion of water and stabilization of the hydrogen-bond network within and between fibril sheets. This mechanism provides a physical basis for a recent experimental observation that electrodeposited chitosan hydrogel film undergoes vertical contraction. The electric field-induced dewetting between amphiphilic chitosan sheets is reminiscent of but fundamentally different from the classic dewetting phenomenon for purely hydrophobic systems, which has been intensively studied by both theoretical and experimental communities in the past. Using active dewetting to control microstructures has implications for tailored engineering of functional materials such as artificial bones and tissues based on self-assembling chitosan.

**TOC Graphic:** 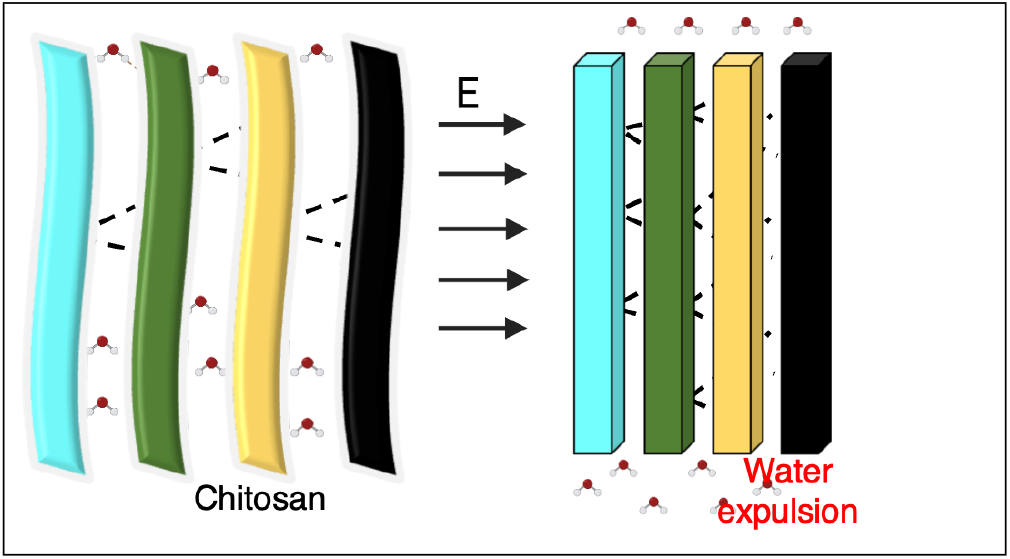

## Introduction

Chitosan is a linear copolymer derived from chitin, made up of D-glucosamine (GlcN) and N-acetylglucosamine (GlcNAc) linked by *β*-1,4-glycosidic bonds. In acidic conditions, the amino groups of chitosan become protonated, making the polymer soluble in aqueous solutions. In neutral or mildly basic conditions (pH above 6.5),^1^ chitosan’s amines deprotonate, leading to self-assembly through hydrophobic effects and hydrogen bonding, forming a three-dimensional hydrogel network with ordered crystalline network junctions.^2^ The pH responsiveness, solubility, tensile properties, and biocompatibility make chitosan a promising material for a wide range of applications, especially in biomedicine and pharmaceutics.^1,3,4^ However, the effective use of chitosan depends on the ability to fabricate it into structures that meet specific biological requirements, e.g., the porous structures that mimic the extracellular matrix of bone.^5^

In recent decades, the use of biomaterials such as chitosan in biomedicine and clinical applications has grown significantly, with even greater expansion anticipated in the coming years.^6^ This surge has highlighted the limitations of traditional fabrication techniques, which typically focus on form and strength to achieve functionality.^5,7^ To address the challenge, new biofabrication methodologies have emerged, with electrobiofabrication at the forefront, which uses electrical signals to control and fabricate material from biological macromolecules such as chitosan, offering the ability to create complex and anisotropic microstructures.^8^ The primary advantage of using electrical signals is the precise control they provide in shaping material behavior, allowing for the development of structures that closely mimic the intricate, hierarchical organization found in biological systems^5,7–9^

During the cathodic electrodeposition of chitosan,^5,8^ mild electrode-imposed voltages drive electrolytic reactions that generate a locally high pH region that deprotonates chitosan’s amines and induces the neutral chitosan chains to self-assemble to form a hydrogel on the electrode surface. These electrical inputs also create an electric field that can exert an electrostatic force to “ recruit” protonated (i.e., charged) chitosan chains from the bulk acidic deposition solution to migrate toward the emergent solution-gel interface.^8^

Despite the apparent simplicity of this electrodeposition mechanism, there are various phenomena occurring at differing length and time scales that control the deposited gel’s supramolecular structure and functional properties. Early work by Liu et al. showed that the addition of salt to the electrodeposition solution dramatically altered the macroscopic structure and mechanical strength of the deposited chitosan gel.^10^ Subsequent work by Yan et al. provided numerous experimental demonstrations suggesting that subtle changes in deposition conditions could yield hydrogels with differing chitosan chain orientations (either perpendicular or parallel to the electrode).^8^ Lei et al. applied this knowledge to create a two-step electrodeposition process (one step with low salt and the second with high salt) to generate a chitosan-based Janus film with dense and porous faces and demonstrated the importance of this structural/functional asymmetry for guided bone regeneration.^11^ In a more recent study, Lei et al. reported chitosan electrodeposition at subambient temperatures and provided experimental evidence that an imposed electric field could also induce structural changes (a contraction) of neutralized chitosan hydrogels.^12^

Despite these advances, detailed knowledge of the molecular processes that underlie chitosan electrodeposition remains extremely limited. The experimental challenge for self-assembly is that phenomena occur at varying length scales and there are not always appropriate tools to measure phenomena at all length scales. All-atom molecular dynamics (MD) simulations can in principle offer molecular details of chitosan’s electrodeposition process under varying conditions such as solution pH, salt concentration, and electric field, although MD simulations of chitosan are very challenging due to the system size (e.g., the degree of polymerization is on the order of 1000). Our earlier work explored the effect of solution salt on the pH-dependent self-assembly of chitosan.^2^ Recently, two MD studies explored the effect of e-field on single chitosan chains in solution. Simulations reported by Yan et al. showed that a charged 20-mer chitosan chain in solution migrates toward the cathode under a constant uniform e-field of 4 mV/nm and a salt concentration of 0.5 M attenuates migration while increasing chain flexibility.^8^ Mahinthichaichan et al. simulated single 5-mer and 10-mer chitosan chains in solution under a constant uniform efield with varying strengths (0, 4, 20 or 400 mV/nm) and found that the large field strength (400 mV/nm) induces significant changes in both conformation and orientation of the chitosan chains.^13^

Although the aforementioned MD studies^8,13^ shed some light on chitosan’s electrodeposition process, questions of how e-field controls the emerging microstructure remain to be addressed. For example, whether and how the e-field directs the chitosan chains to self-assemble in a specific orientation^8^ and why contraction occurs to the hydrogel after gelation?^12^ In this work, we conducted all-atom MD simulations to investigate the effect of a constant uniform e-field on the microstructure of chitosan’s crystalline region. As a model of the crystalline region, the simulations started from a chitosan nanofibril comprised of 4 antiparallel sheets, each with 6 parallel 20-mer chains based on the X-ray diffraction structure of anhydrous chitosan crystals.^14^ The simulations show that under a mild e-field of 4 mV/nm, the chitosan nanofibril becomes more ordered and contracts in the sheet stacking distance. Moreover, the chitosan nanofibril comprised of 4 sheets has no preferred orientation in the e-field. The analysis reveals that while the nanofibril becomes hydrated in the field-free simulations, the application of an e-field induces the expulsion of water from the fibril while stabilizing the direct hydrogen bond network. This e-field induced dewetting transition is reminiscent of but differs from the passive dewetting phenomenon extensively studied in the past. Using active dewetting to control microstructures has implications for tailored engineering of functional materials such as artificial bones and tissues based on self-assembling chitosan.

## Experimental

The model chitosan nanofibril studied in this work (Fig. 1) was built from the asymmetric unit of the X-ray structure of the anhydrous chitosan crystals at 1.17 Å resolution^14^ using the program Mercury.^15^ The unit cell is in the *P* 2_1_2_1_2_1_ space group, with *a* = 8.129, *b* = 8.347, *c* = 10.311 Å . The unit cell contains three chains (1 dimer and 2 monomers). According to the authors,^14^ the molecular arrangements in the parallel, hydrogen-bonded sheets are very similar to cellulose II along ac plane and cellulose III_I_ along bc plane. The anhydrous chitosan was prepared by heating the hydrated chitosan to 200°C.^14^ To reduce computational costs, we constructed the smallest nanofibril that maintained structural stability in our previous simulations,^2^ composed of 4 antiparallel sheets, each containing 6 chains of 20 neutral glucosamine units. At the start of the simulations, the nanofibril was oriented such that the directions of sheet growth, sheet stacking, and fibril elongation aligned with the x-, y-, and z-axes, respectively (Fig. 1d). The dimensions of the nanofibril were 46.7 Å along the sheet growth direction, 12.1 Å along the sheet stacking direction, and 103.1 Å along the fibril elongation (Fig. 1d).

**Figure 1.**
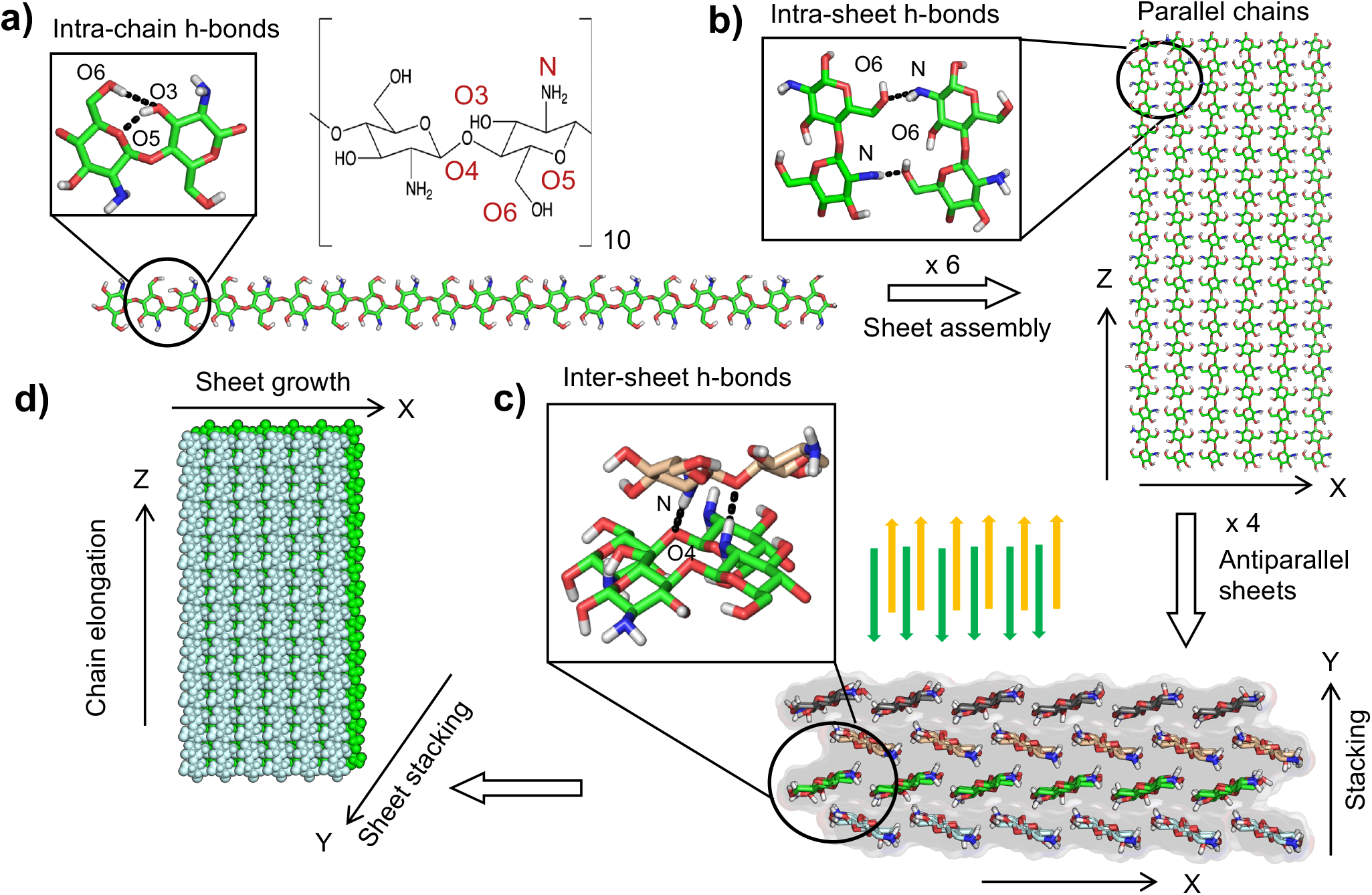
Schematic of the model chitosan nanofibril built from the X-ray diffraction structure of anhydrous chitosan. ^**14**^. **a)** A chitosan chain made up of 20 glucosamine units. Adjacent glucosamine units form intra-chain H-bonds, O3^*′*^H-O5 and O6H–O3^*′*^. **b)** A nanofibril sheet made up of 6 parallel chitosan chains, stabilized by the N^*′*^H– O6 intra-sheet H-bond. **c)** A chitosan nanofibril comprised of 4 anti-parallel stacked sheets. Sheet-sheet stacking is stabilized by the inter-sheet N^*′*^H–O4 H-bond and hydrophobic contacts. **d)** At the start of the simulations, the fibril sheets are on the xz plane, with the chitosan chains elongate along z, while the sheets are stacked along y.

All-atom molecular dynamics (MD) simulations were performed using the AMBER20 program.^16^ Chitosan was represented by the modified^2,17^ CHARMM36 carbohydrate force field.^18,19^ Water was represented by the CHARMM-style^20^ TIP3P model.^21^ CHAMBER^22^ was used to convert the CHARMM parameter and topology files to the AMBER format. The nanofibril was solvated in a 120 Å x 120 Å x 120 Å cubic water box, resulting in a system size of approximately 266,667 atoms. The solvated system underwent energy minimization with a harmonic restraint potential (force constant of 10 kcal/mol/Å ^2^) placed on the heavy atoms. The system was minimized for a total of 4000 steps. The first 1000 steps were conducted with the steepest descent algorithm followed by 3000 steps using the conjugate gradient algorithm. The system was then heated over 10 ns to 300 K using a time step of 2 ps in the NVT ensemble. Afterwards, the system was equilibrated for a total of 100 ns, in which the harmonic potential force constant was gradually reduced from 5, 2.5, 1, 0.1 to 0 kcal/mol/Å ^2^ under constant NPT conditions. The temperature was maintained using a Langevin thermostat,^23^ and the pressure was maintained at 1 bar using the Monte Carlo barostat.^24^ Note, e-field was absent in energy minimization, heating, and equilibration.

Following the equilibration, a total of six independent production simulation runs were conducted. Three production runs were performed for the field-free control system, while one production run was performed for each system in which a constant uniform e-field of 4 mV/nm was applied in x, y, or z direction. The van der Waals interactions were smoothly switched to zero from 10 to 12 Å . The particle mesh Ewald method^25^ was used to calculate long-range electrostatic energies with a sixth-order interpolation and 1 Å grid spacing. Periodic boundary conditions were applied throughout the simulations. Bonds involving hydrogens were constrained using the SHAKE algorithm^26^ to enable a 2-fs timestep. All other details can be found in the freely downloadable input files (see Data Availability). Simulation analysis was performed using CPPTRAJ^27^ and VMD.^28^ Unless otherwise noted, the last 500 ns data was used for analysis.

Volume of the nanofibril was calculated using the Qhull program in the Python library SciPy,^29^ which defines volume as the convex hull *Conv*(*P*). Given a set of points

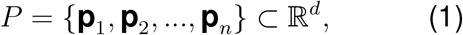

*Conv*(*P*) is defined as

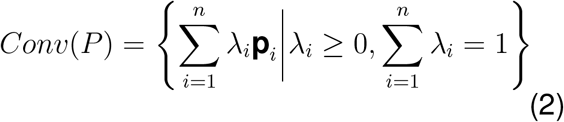

The convex hull is tessellated into non-overlapping simplices and the total volume is calculated as the sum of the volumes of each simplex. Each simplex volume in R^*d*^ formed by *d* + 1 points is given by the following formalism:

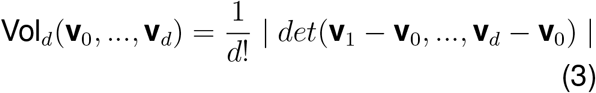

The volume change, ΔVolume is calculated as

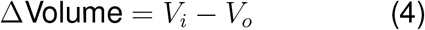

where V_*o*_ is the volume of the fibril at the start of the simulation and V_*i*_ is the volume of the fibril at a chosen time point in the simulation.

## Results and Discussion

### H-bond network provides structural foundation of the chitosan nanofibril

To construct a model chitosan nanofibril, we utilized the high-resolution X-ray fiber diffraction structure of anhydrous, fully deacetylated chitosan in the *β* allomorph.^14^ While an earlier study reported an X-ray structure of *α*-chitosan,^30^ the *β*-form X-ray structure was selected due to its superior resolution (1.17 Å)^14^ . Note, the choice of allomorph does not influence the investigation of the e-field effects. For computational feasibility, the model chitosan nanofibril contains 24 chitosan chains, each having 20 glucosamine units (Fig. 1a). The chitosan chains are elongated along the z-axis, while 6 parallel chains form a sheet on the xz plane (Fig. 1b) and 4 antiparallel sheets are stacked along the y-axis to form the nanofibril (Fig. 1c and d).

The chitosan nanofibril is stabilized by intrachain, intra-sheet, and inter-sheet hydrogen bonds (H-bonds) as well as hydrophobic contacts. Based on the H-bond donor-acceptor heavy-atom distance, the dominant intrachain H-bonds are O3^*′*^H–O5 and O6H–O3^*′*^ (Fig. 1a) and the dominant intra-sheet H-bond N^*′*^H–O6 is found between the parallel chains while the inter-sheet N^*′*^H–O4 H-bond is found between the antiparallel sheets (Fig. 1c).

A total of six simulations were conducted, three in the absence (1 *µ*s production run each) and three in the presence of a constant uniform e-field of 4 mV/nm along x, y or z direction (6 *µ*s production run each). Prior to the production runs, the field-free equilibration run was conducted for 100 ns. Here water entered the fibril forming H-bonds with N, O6, O3 and O5 atoms. Interestingly, water molecules also participated in bridging the inter- and intra-sheet H-bonds primarily between N and O6 atoms (see later discussion). The simulations were conducted until the properties of interest reached convergence (Supplemental Fig. S1–S4). Unless otherwise noted, data from the last 500 ns was used for analysis. Note, in order to observe potential effects of e-field within microsecond simulation time, which is roughly 7 orders of magnitude shorter than the experimental timescale of minutes, we applied an e-field that is 1000 times greater in magnitude than the experiment^8^ but nonetheless comparable to that (*∼* 10 mV/nm) naturally imposed across biological membranes.^31,32^

### E-field stabilizes intra-chain H-bonding and increases order of the nanofibril

Consistent with our previous simulation work,^2,33^ the intra-chain O3^*′*^–O5 H-bond is dominant in all simulations (occupancies greater than 74%), while the formation of the O6H–O3^*′*^ H-bond suggested by the fiber diffraction data is minimal with the occupancies below 10% (Fig. 2a). Surprisingly, in the presence of the e-field regardless of its direction, the O3^*′*^–O5 H-bond is stabilized, as demonstrated by the 6% increase in the H-bond occupancy (Fig. 2a). Since the O3^*′*^– O5 H-bonds stabilize a chitosan chain in the *syn* conformation which corresponds to a larger glycosidic linkage angle,^33^ the chitosan chains in the nanofibril become more extended, as shown by the sharp peak in the end-to-end distance distribution from the simulations with e-field (Fig. 2b). Interestingly, along with increased chain rigidity, the nanofibril becomes more ordered in the efield, as demonstrated by the decrease in the root mean square fluctuation (RMSF) of chain atoms around their average positions (Fig. 2c). Indeed, visualization of the trajectory snapshots confirms that the nanofibril is flexible in the absence of field and becomes more ordered in the presence of an e-field (Fig. 2d).

**Figure 2.**
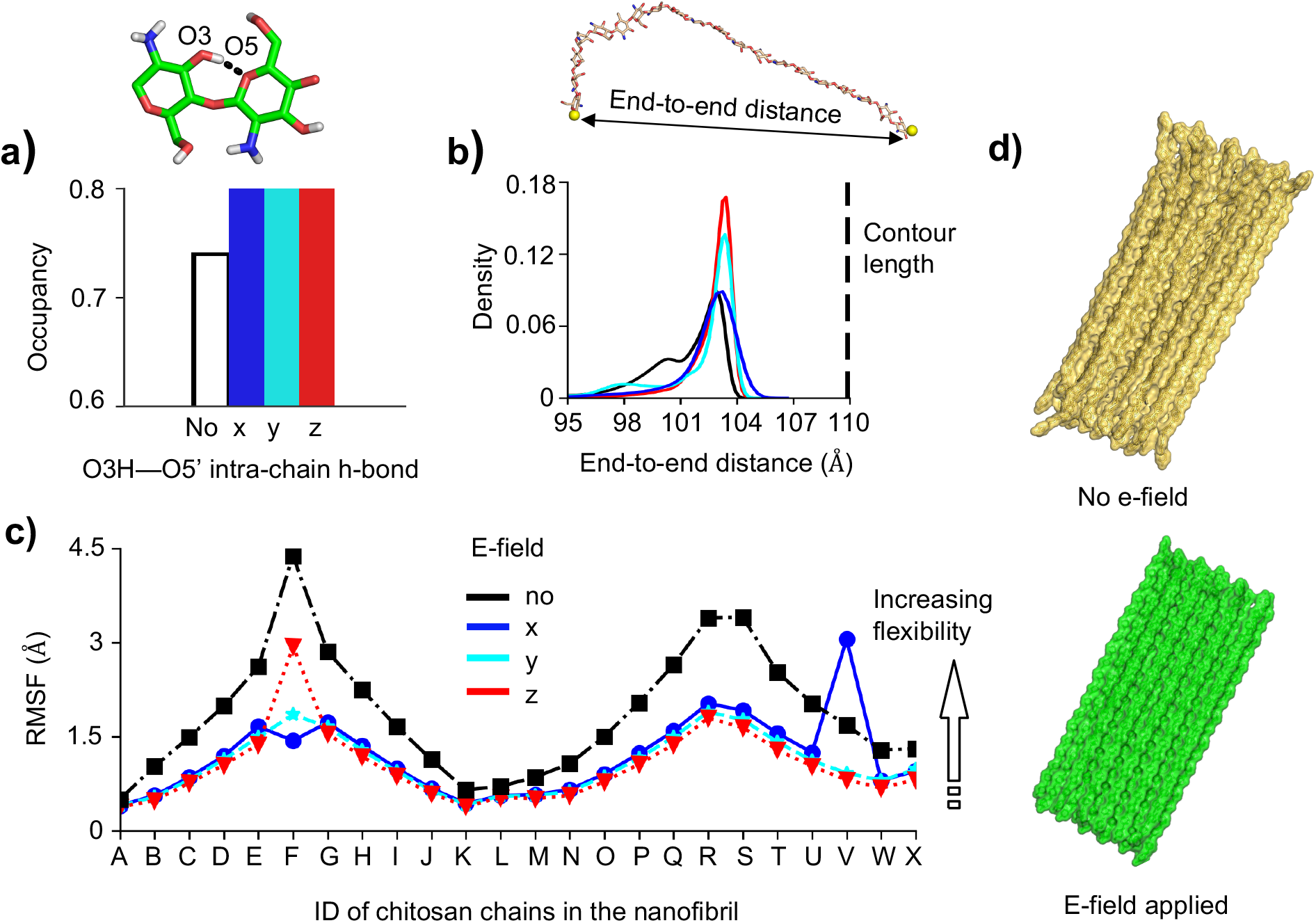
E-field stabilizes intra-chain H-bonding, making the fibril sheets more ordered. **a)** Occupancy of the O3H–O5’ intra-chain H-bond (snapshot above) for all chains in the nanofibril without (black) and with an e-field (blue, cyan, and red). **b)** Probability distribution of the chitosan chain end-to-end distance, defined as between O4 atoms of glucosamine unit 1 and unit 20. The fully stretch-out (contour) length is shown. Calculations used all chains in the nanofibril. **c)** Heavy-atom root mean square fluctuation (RMSF) of the chitosan chains in the nanofibril without (black) and with e-field (blue, cyan, and red). **d)** Representative snapshots of the nanofibril from the field-free simulation run 3 (top) and from the simulation with the e-field applied in the z-direction (bottom).

### E-field stabilizes inter-chain H-bonds while destabilizing chitosan-water H-bonds

Now we examine whether and how the e-field impacts the inter-chain (i.e., inter- and intrasheet) H-bonding in the chitosan nanofibril. Specifically, we calculated the occupancies of all possible inter- and intra-sheet H-bonds in the absence and presence of the e-field (Fig. 3a and b). Surprisingly, both the dominant inter-sheet NH–O4 and intra-sheet O6H– N H-bonds are stabilized in simulations with an e-field in any direction, as evident from the increased occupancies relative to the field-free simulations (Fig. 3a). Note that NH– O4 H-bonds stabilize sheet-sheet stacking along with the hydrophobic contacts, while the O6H–N H-bond which is the strongest H-bond based on the diffraction data^14^ stabilizes the self-assembled sheet. It is also interesting to note that while the NH–O6 H-bond is present in the initial nanofibril structure, it is infrequently formed (occupancy about 10%) in all simulations.

**Figure 3.**
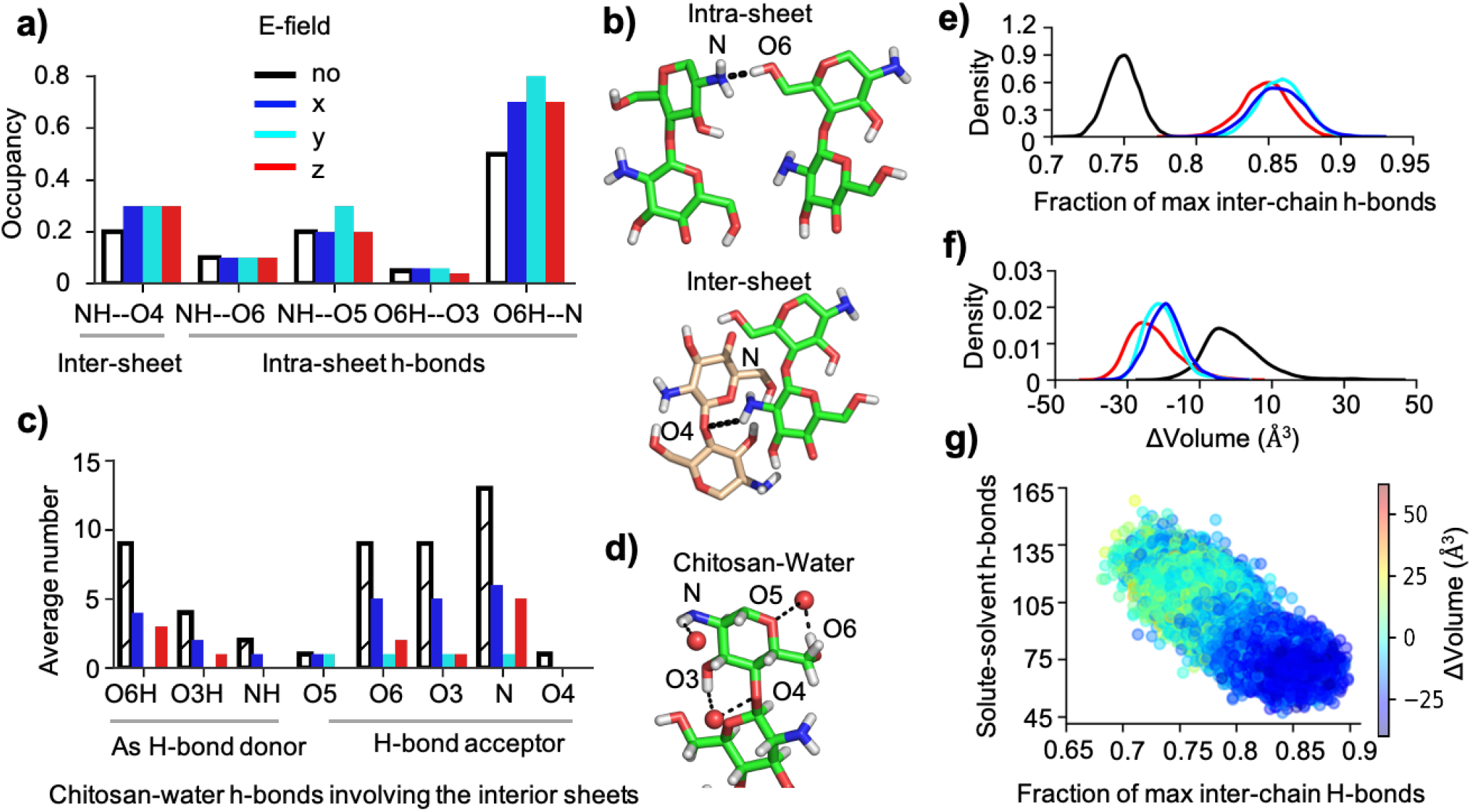
E-field stabilizes intra- and inter-sheet H-bonding while destabilizing the chitosan-water H-bonds, leading to fibril’s volume contraction. **a)** Occupancies of specific inter-sheet and intra-sheet H-bonds in the chitosan nanofibril without and with e-field applied in the x-(blue), y-(cyan), or z-direction (red). An H-bond is considered form if a donor-acceptor heavy-atom distance is below 3.5 Å and the donor-H-acceptor angle is greater than 135^*°*^. **b)** Representative snapshots from the field-free simulation (run 3) showing the most abundant intra-sheet and inter-sheet H-bonds. **c)** Average number of the specific chitosan-water H-bonds involving the (12) chains in the (2) fibril inner sheets as a H-bond donor (left) or acceptor (right). **d)** Representative snapshots from the field-free simulation (run 3) showing chitosan-water H-bonds. **e**,**f)** Probability distribution of the fraction of max inter-chain (inter- and intra-sheet) H-bonds and volume change (ΔVolume) of the chitosan nanofibril without (black) and with an e-field (colored). Fraction of max H-bonds is calculated relative to the total number of H-bonds in the energy minimized structure of the chitosan nanofibril. ΔVolume is relative to the energy minimized state, calculated by the Qhull library of SciPy ^34^ using the convex hull algorithm. ^35^ The field-free analysis is based on 3 independent runs. **g)** Total number of chitosan-water H-bonds formed by the 12 chains of the interior sheets vs. the fraction of max inter-chain H-bonds from the simulation with an e-field applied in the z-direction. To clearly show the correlation, data from the entire simulation is used. Data is color-coded according to ΔVolume of the nanofibril. Blue and red indicates contraction and expansion, respectively.

We hypothesized that stabilization of intra- and inter-sheet H-bonding may be related to destabilization of chitosan-water interactions. To test it, we calculated the occupancies of all possible chitosan-water H-bonds formed by the two interior fibril sheets (Fig. 3c and d). The two exterior sheets were excluded from this analysis as they have a solvent-facing side. Indeed, the number of the major chitosan-water H-bonds, involving O6, O3, and N, is reduced when an e-field is applied in any direction (Fig. 3c). This confirms that chitosan-water and chitosan-chitosan H-bonds are in direct competition. For example, the decrease in the number of O6H–water and N—water H-bonds (Fig. 3c,d) can be explained by the higher occupancy of the intrasheet O6H–N H-bond (Fig. 3a,b).

### Increased inter-chain and decreased chitosan-water H-bonding is correlated with fibril’s volume contraction

Consistent with the increased occupancies of the individual inter-chain H-bonds under an efield (Fig. 3a), the total number of inter-chain H-bonds increases by about 13% in the presence of an e-field regardless of the direction (Fig. 3e). Interestingly, the fibril’s volume decreases under the e-field regardless of its direction, as shown by the left shift of the ΔVolume distributions by about 20–25 Å ^3^ from the field-free simulations (Fig. 3f). In the latter case, the ΔVolume distribution is centered around 0 Å ^3^, demonstrating that on average no volume change is observed in the absence of an e-field. To test whether volume contraction is related to H-bonding, a scatter plot of the number of chitosan-water H-bonds involving the 12 interior chains vs. the fraction of maximal inter-chain H-bonds was made, whereby the data points are color-coded by fibrils’ volume change (Fig. 3g). It is evident that as the number of inter-chain H-bonds increases, the number of chitosan-water H-bonds as well as the volume of the nanofibril decreases (negative ΔVolume, Fig. 3g).

### Decreased hydration of fibril interior leads to compaction of sheet-sheet stacking

Formation of less chitosan-water H-bonds by the interior chains (Fig. 3g) suggests that the interior of chitosan is less hydrated. The distribution of the number of water in the first hydration shell confirms the loss of water under an applied e-field (Fig. 4a). Next, we specifically examined the water between two interior sheets (Fig. 4b and 4d). Remarkably, e-field induces a significant reduction in the number of interior water. Moreover, the extremely broad distribution observed in the absence of e-field contrasts sharply with the peaked distributions seen under the e-field, suggesting that water molecules in exchange with the bulk, i.e., the unbound water not in H-bonding with chitosan chains, are removed (Fig. 4b).

**Figure 4.**
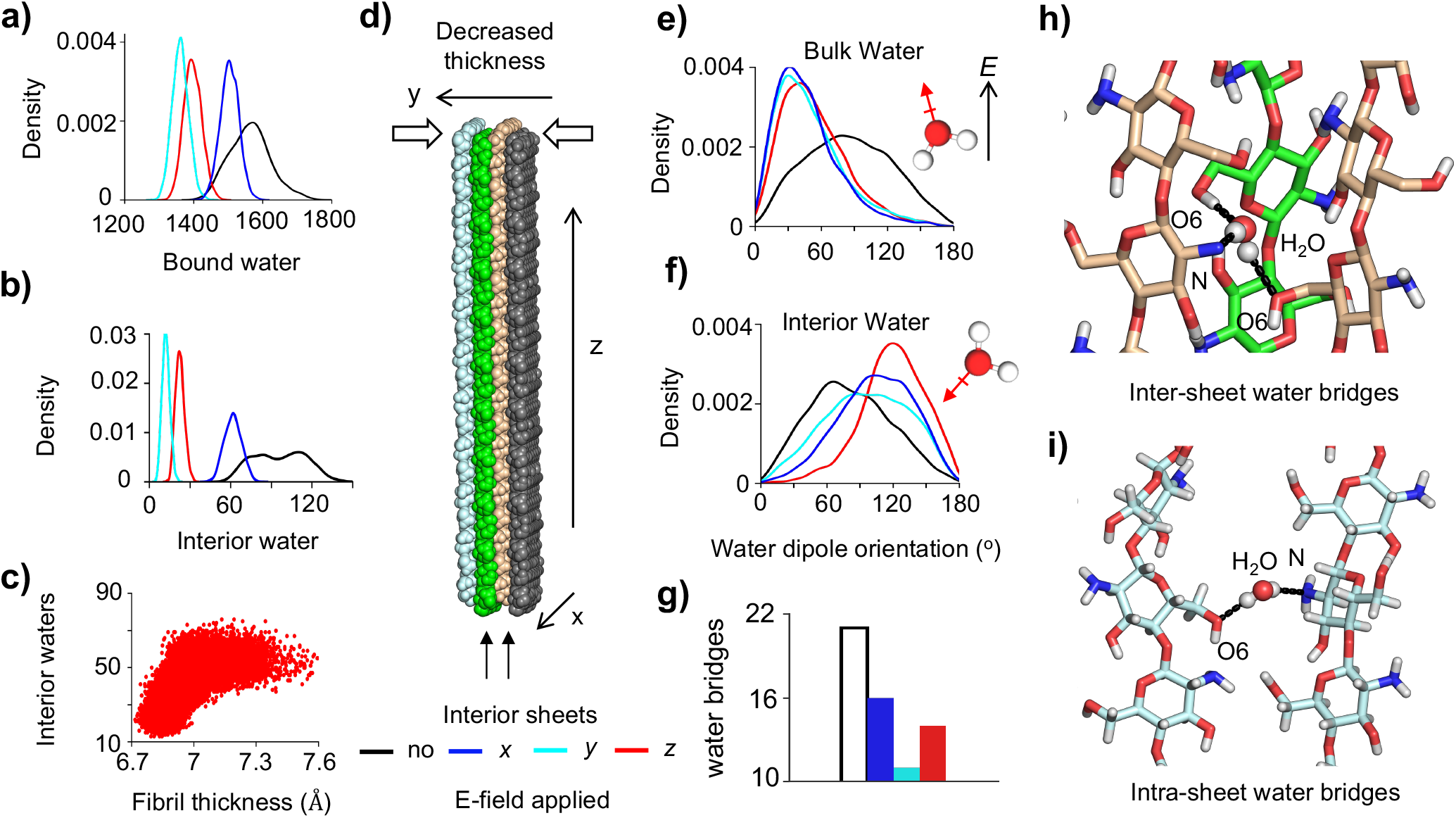
Loss of interior water, decreased fibril thickness, and reduced number of water bridges resulted from the field-dipole interactions. **a)** Probability distribution of the number of water bound to the chitosan nanofibril without (black) and with e-field (colored). Bound water refers to any water whose oxygen is within 3.5 *Å* radius from any heavy atom of chitosan. **b)** Number of water between the two interior sheets. **c)** Scatter plot showing the number of interior water is correlated with the inter-sheet distance defined as between the center of mass of the middle chains in sheet 2 and 3. The trajectory with e-field applied in the z direction is used. **d)** A schematic showing decreased fibril thickness. **e, f)** Distributions of the dipole orientations of the bulk water (f) and the bound water between two interior sheets (g). The e-field direction is used as a reference. Dipole orientations without the e-field are in reference to the z-axis. Bulk water is defined as those whose oxygen atom is at least 10 Å away from any heavy atom of chitosan. **g)** Number of water-bridged H-bonds between the two interior sheets. **h, i)** Snapshots showing the inter(top) and intra-sheet (bottom) water-bridged H-bonds. Light blue, green and wheat colors indicate different fibril sheets.

We hypothesized that the reduction in interior water results in decreased fibril thickness (i.e., compaction of sheet-sheet stacking), which may explain the decreased fibril volume (Fig. 3f). Although a shrinkage of sheet width (i.e., intra-sheet chains are closer to each other) may also decrease fibril’s volume, this is highly unlikely, as the chitosan chains within a fibril sheet are stably assembled by strong intra-sheet H-bonds between glucosamine units of adjacent chains, particularly the H-bond interactions between the hydroxyl O6H and the amino nitrogen (Fig. 1b). Indeed, the distribution plots of fibril width demonstrate that the applied e-field rigidifies the sheet structure and does not decrease its dimension (SI Figure S6). In contrast, the sheet-sheet stacking distance is correlated with the number of interior water and is decreased by 0.5–0.7 Å with an applied e-field (Fig. 4c).

### Disruption of water bridges is due to fielddipole interactions

According to classical electrodynamics, a torque is exerted on a dipole under the influence of a uniform e-field. The torque, whose magnitude is proportional to the dipole moment, rotates the dipole until the dipole is aligned with the e-field. We hypothesized that the observed fibril dehydration is due to the e-field interaction with the water dipoles so that the ideal chitosan-water H-bond geometries cannot be satisfied. Note, polarizability of water is not considered in the present MD simulations.

To test the above hypothesis, we examined the dipole orientations of bulk water molecules and those bound to the two interior sheets (interior water). Without an e-field, water is expected to freely rotate in the bulk, which is indeed the case, as evident from the nearly flat distribution of the dipole orientation angle with respect to the z axis (Fig. 4e, black curve). Under an applied e-field, the bulk water dipoles are reoriented to a more or less parallel alignment (about 30^*°*^) with respect to the field direction (Fig. 4e, colored curves), consistent with the aforementioned prediction from physics. Note, the e-field does not affect the diffusion of water molecules, as the mean square displacement is identical for all simulations in the presence or absence of e-field. However, interior bound water molecules cannot freely rotate due to the constraints of Hbonding with the chitosan chains (primarily in the form of water bridges). Thus, the e-field interaction with water dipoles disrupts many water-chitosan H-bonds (Fig. 4g), expelling these water molecules to the bulk phase while leaving only a few strong water-bridged interactions intact (sharp peaks in Fig. 4b). These remaining interactions exhibit preferential water dipole orientations that differ from those observed in the absence of an e-field (Fig. 4f). Another consequence of disruption of water bridges (Fig. 4g) is the replacement of water-mediated H-bonds with direct H-bonds (Fig. 3a), which results in a decrease in the sheet-sheet stacking distance. Lastly, a similar disruptive effect is observed for intra-sheet water bridges (Fig. 4i) under an e-field. However, due to the extensive network of intrasheet H-bonds (Fig. 3a), the replacement of some water bridges with direct H-bonds increases the structural order without altering the dimension of the fibril sheets (SI Fig. S6).

### Chitosan nanofibril has no preferred orientation in e-field due to dipole cancellation

Considering that the e-field induces reorientation of water molecules, we asked if it can also have an orientational effect on the chitosan fibril. To address this question, we first examined the dipole of a single chitosan chain. For a fully extended 20-mer chitosan chain, the dipole is nearly aligned with the chain vector, with an orientation angle of *∼*6^*°*^. Here, the chain vector is defined as extending from the center of mass of the second to the 19th sugar unit.

In the absence of an e-field, the chitosan chains in the nanofibril are flexible and adopt many different conformations (Fig. 2b and c). As a result, the chain dipole is no longer aligned with the chain and instead samples an angle around 65^*°*^ between 40^*°*^ and 90^*°*^ (Fig. 5a, top). In contrast, under the e-field, the chains are rigid and resembles the ideal extended state (Fig. 2b and c). Consequently, the chain dipoles are nearly parallel to the chain vector, with the most probable angle of *∼* 10^*°*^ (Fig. 5a, bottom).

**Figure 5.**
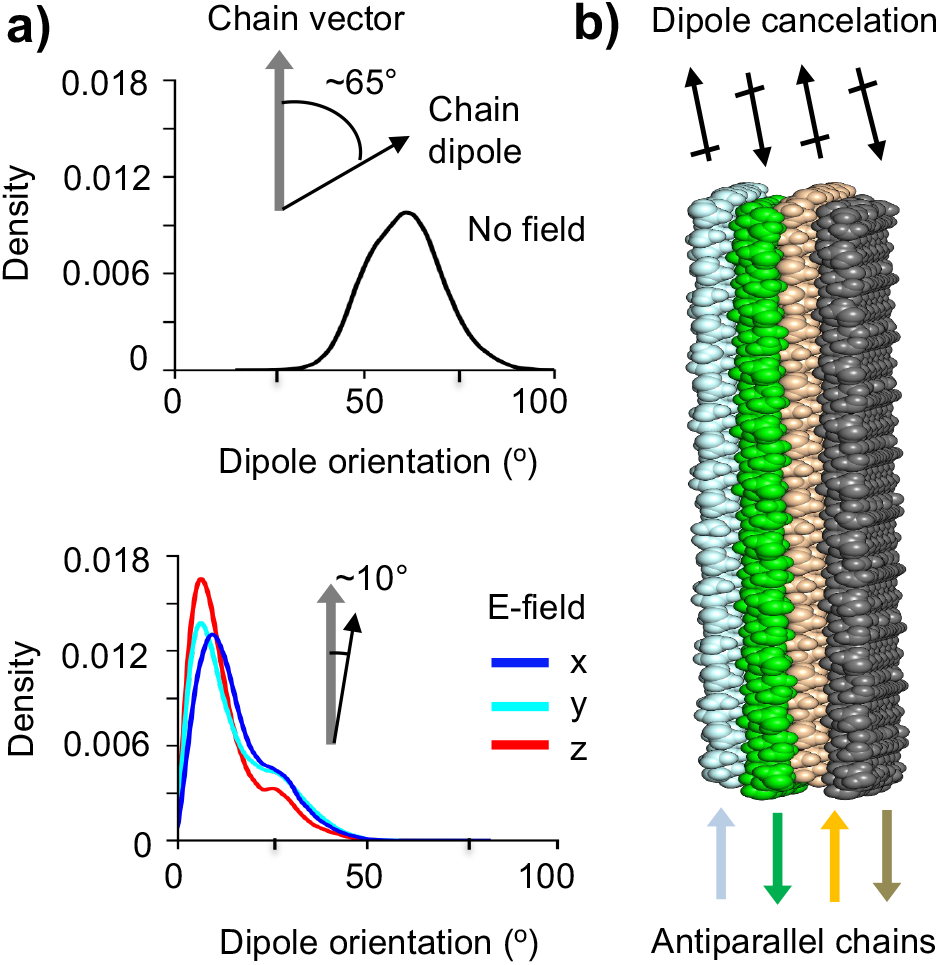
The chitosan chain dipoles in the nanofibril are nearly aligned with the direction of chain elongation. **a)** Probability distributions of the orientation angle of the chitosan chain dipole (black arrows) with respect to the chain vector (grey arrow) in the simulations without (top) and with an e-field in the x, y, or z direction (bottom). **b)** Illustration of the cancellation of chitosan chain dipoles (black arrows) in the nanofibril made up of 4 antiparallel sheets. Each sheet is comprised of parallel chitosan chains while adjacent sheets have antiparallel chains. Colored arrows indicate the chain vector directions in different sheets.

Since the chitosan chain dipoles are aligned nearly parallel to the chain, an applied electric field would orient the nanofibril such that the chain elongation axis points toward the direction of the e-field. However, this is not the case. In all three simulations with an efield, the chitosan nanofibril rotates in solution without a preferred orientation, similar to the simulations without an e-field. This can be explained by the antiparallel arrangement of chitosan chains in the adjacent sheets, which leads to dipole cancellation. Since there are four sheets, the net dipole moment of the nanofibril is zero (Fig. 5b), which explains why the nanofibril does not have a preferred orientation in e-field. We note that the same applies to *α*-chitosan, where intra-sheet chains are antiparallel to each other, resulting in a total net zero dipole moment.

## Conclusion

Our MD simulations offer an atomic-level view of how an applied e-field impacts the self-assembled structure of chitosan in solution. The simulations started with an anhydrous nanofibril (based on the X-ray diffraction structure^14^) comprising 4 antiparallel stacked sheets, each having 6 parallel chitosan chains. In the absence of e-field, the nanofibril undergoes hydration, forming water-mediated intra- and inter-sheet H-bonds, consistent with the X-ray diffraction data of hydrated chitosan.^36^ The nanofibril exhibits significant dynamics, characterized by flexible chitosan chains and fluctuating sheet dimensions. When subjected to a mild e-field (4 mV/nm), the nanofibril shows enhanced structural organization while simultaneously contracting along the sheet stacking axis. This behavior is consistent with recent electrodeposition experiments, which demonstrated that neutralized chitosan hydrogels contract in the direction perpendicular to the electrode under a uniform e-field^12^ (Fig. 6b). Note, although the specific allomorph formed by the self-assembled chitosan under the e-field is unknown and may differ from the one studied by our simulations, the above conclusions remain unchanged because the observed dewetting transitions are independent of chitosan chain alignment directions. The simulations also showed that the chitosan nanofibril adopts random orientations under the e-field, which is attributed to the net zero dipole moment. This conclusion also applies to *α*-chitosan, which has a net zero dipole moment due to antiparallel chains.

**Figure 6.**
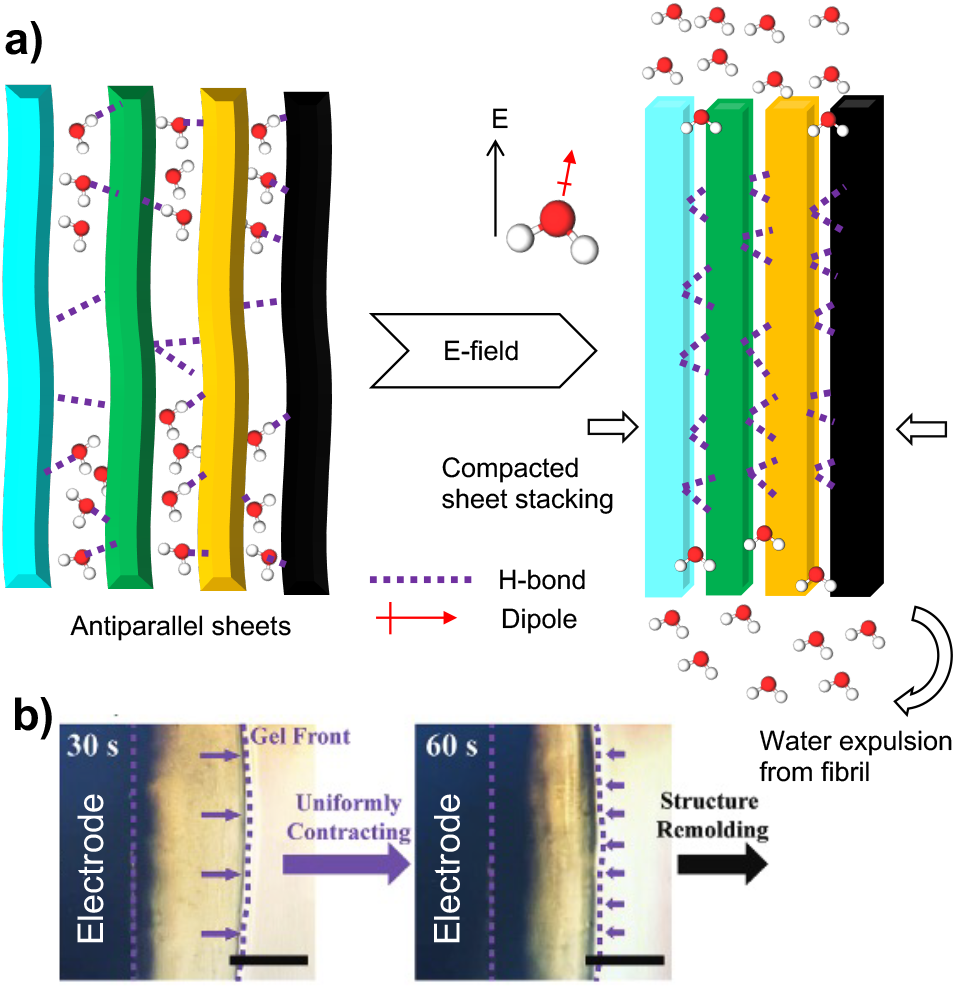
A working model of how e-field remodels the microstructure of chitosan hydrogel’s crystalline regions through dewetting. **a)** Application of an e-field in any direction reorients water molecules in the bulk and disrupts water-bridged inter- and intra-sheet H-bonds in the hydrogel’s crystalline regions. The latter causes water molecules to be driven out of the fibril structure, i.e., dewetting, leading to fibril contraction along the sheet-stacking direction and increased order as a result of a more dense network of inter- and intra-sheet H-bonding. **b)** An electrodeposition experiment ^12^ shows the contraction of neutralized chitosan hydrogels deposited on the electrode under a uniform e-field. The image is reproduced from the open-access article by Lei et al ^12^ under a Creative Commons CC BY license.

The analysis of simulation data demonstrated that fibril contraction and increased order is the result of a stabilized H-bonding network, which includes the intra-sheet H-bonds that stabilize the fibril sheets as well as the inter-sheet H-bonds that stabilize and compact the sheet stacking (Fig. 6a). Our analysis further revealed that the effects of the e-field are mediated by solvent through interactions between field and water dipoles. Through field-dipole interactions, water-mediated Hthan 1 nm^2^)^41^ hydrophobic plates approaching a critical distance (*D*_*c*_), creating a thin vapor layer which leads to strong attractive forces and subsequent hydrophobic collapse.^37–40,44^ Our recent simulations of the self-assembly process of chitin (acetylated chitosan)^45^ demonstrated a signature of hydrophobic effect,^39,40^ i.e., an increase in temperature enhances hydrophobic forces; however, the lubricating role of water as previously discussed in protein folding simulations^46^ was also observed, in which inter-chain H-bond formation is concomitant with the expulsion of water in the final stage of self-assembly. The lubricating role of water was also demonstrated in an early simulation study of chitosan’s self-assembly.^2^ On the other hand, bonds are disrupted, leading to the sequestration of water from the fibril, which in turn stabilizes the direct H-bond network, reducing the inter-sheet distances and making the fibril more ordered (Fig. 6a). Note that the dipoles discussed thus far refer to permanent dipoles arising from partial atomic charges as defined by classical force fields. Induced dipoles under the mild electric field applied here are expected to be negligible and are not anticipated to influence the conclusions of this work.

The finding that the e-field induces dehydration and compaction of the chitosan fibril is reminiscent of the dewetting phenomenon extensively discussed in the theoretical literature.^37–43^ Accordingly, dewetting or drying occurs between two sufficiently large (greater the hydrophobic nature of chitosan interior is supported by the p*K* _a_ downshifts of the amino groups.^2,47^

The aforementioned data suggests that chitosan’s fibril sheets exhibit an amphiphilic nature: the extensive sugar backbones create hydrophobic regions, while polar nitrogen and oxygen atoms form readily accessible H-bond sites that can interact with water molecules. Without an applied e-field, water can enter, forming water-chitosan and water-bridged inter- and intra-sheet H-bonds; however, with an applied e-field, the water bridges are disrupted, restoring the direct inter- and intra-sheet H-bond network. Therefore, the e-field effectively weakens the hydrophilic nature of chitosan’s fibril sheets, enabling dewetting transitions typical for hydrophobic plates that have been intensively studied by theorists^37–43^ and later confirmed in purely hydrophobic nanoporous systems by experimentalists.^48,49^ Chitosan’s amphiphilic character creates a distinct e-field-induced dewetting mechanism that fundamentally differs from the classic wetting and dewetting behavior of hydrophobic systems.

Our finding suggests that e-field can serve as an effective stimulus for programming both the hydration and fibrillar structure of self-assembling materials. Considering that chitosan’s dewetting transitions were observed using an e-field of comparable magnitude to those across the biological membranes and amphiphilic systems are common in biology, our findings have implications for understanding cellular processes. It is intriguing to consider whether biology uses e-fields to help create the hierarchical organization and functional properties of structural polymers, e.g., cellulose in cell walls and protein collagen in tissues. Active dewetting control of microstructures has implications for tailored engineering of functional materials such as artificial bones and tissues based on self-assembling chitosan.

## Supporting Information

Supplemental figures display the time series of the calculated number of hydrogen bonds, solvent accessibility, nanofibril volume and thickness for all simulations. An additional supplemental figure shows the probability distribution of the nanofibril dimension along the sheet growth direction for all simulations.

## Data Availability

All simulation inputs and data for plotting are freely downloadable at https://github.com/ JanaShenLab/Chitosan_efield/

## Acknowledgments

We thank a former PhD student Kevin Tsai for his unpublished MD simulations of a single chitosan chain in an applied e-field. Financial support from National Science Foundation (CBET1932963) and National Institutes of Health (R35GM148261) is acknowledged.

## References

(1) Rinaudo, M. Chitin and Chitosan: Properties and Applcations. Progress in Polymer Science. Prog. Polym. Sci. 2006, 31, 603–632.

(2) Morrow, B. H.; Payne, G. F.; Shen, J. pH-Responsive Self-Assembly of Polysaccharide through a Rugged Energy Landscape. J. Am. Chem. Soc. 2015, 137, 13024–13030.

(3) Kumar, M. N. V. R.; Muzzarelli, R. A. A.; Muzzarelli, C.; Sashiwa, H.; Domb, A. J. Chitosan Chemistry and Pharmaceutical Perspectives. Chem. Rev. 2004, 104, 6017–6084.

(4) Jimenez-Gomez, C. P.; Cecilia, J. A. Chitosan: A Natural Biopolymer with a Wide and Varied Range of Applications. Molecules 2020, 25, 1–43.

(5) Li, J.; Wu, S.; Kim, E.; Yan, K.; Liu, H.; Liu, C.; Dong, H.; Qu, X.; Shi, X.; Shen, J.; Bentley, W. E.; Payne, G. F. Electrobiofabrication: electrically based fabrication with biologically derived materials. Biofabrication 2019, 11, 032002.

(6) Lele, M.; Kapur, S.; Hargett, S.; Sureshbabu, N. M.; Gaharwar, A. K. Global trends in clinical trials involving engineered biomaterials. Sci. Adv. 2024, 10, eabq0997.

(7) Ling, S.; Kaplan, D. L.; Buehler, M. J. Nanofibrils in Nature and Materials Engineering. Nat. Rev. Mater. 2018, 3, 18016.

(8) Yan, K.; Liu, Y.; Zhang, J.; Correa, S. O.; Shang, W.; Tsai, C.-C.; Bentley, W. E.; Shen, J.; Scarcelli, G.; Raub, C. B.; Shi, X.-W.; Payne, G. F. Electrical Programming of Soft Matter: Using Temporally Varying Electrical Inputs To Spatially Control Self Assembly. Biomacromolecules 2018, 19, 364–373.

(9) Nepal, D.; Kang, S.; Adstedt, K. M.; Kanhaiya, K.; Bockstaller, M. R.; Brinson, L. C.; Buehler, M. J.; Coveney, P. V.; Dayal, K.; El-Awady, J. A.; Henderson, L. C.; Kaplan, D. L.; Keten, S.; Kotov, N. A.; Schatz, G. C.; Vignolini, S.; Vollrath, F.; Wang, Y.; Yakobson, B. I.; Tsukruk, V. V.; Heinz, H. Hierarchically Structured Bioinspired Nanocomposites. Nat. Mater. 2023, 22, 18–35.

(10) Liu, Y.; Zhang, B.; Gray, K. M.; Cheng, Y.; Kim, E.; Rubloff, G. W.; Bentley, W. E.; Wang, Q.; Payne, G. F. Electrodeposition of a Weak Polyelectrolyte Hydrogel: Remarkable Effects of Salt on Kinetics, Structure and Properties. Soft Matter 2013, 9, 2703.

(11) Lei, M.; Qu, X.; Liu, H.; Liu, Y.; Wang, S.; Wu, S.; Bentley, W. E.; Payne, G. F.; Liu, C. Programmable Electrofabrication of Porous Janus Films with Tunable Janus Balance for Anisotropic Cell Guidance and Tissue Regeneration. Adv. Funct. Mater. 2019, 29, 1900065.

(12) Lei, M.; Liao, H.; Wang, S.; Zhou, H.; Zhao, Z.; Payne, G. F.; Qu, X.; Liu, C. Single Step Assembly of Janus Porous Biomaterial by Sub-Ambient Temperature Electrodeposition. Small 2022, 18, 2204837.

(13) Mahinthichaichan, P.; Tsai, C.-C.; Payne, G. F.; Shen, J. Polyelectrolyte in Electric Field: Disparate Conformational Behavior along an Aminopolysaccharide Chain. ACS Omega 2020, 5, 12016–12026.

(14) Naito, P.-K.; Ogawa, Y.; Sawada, D.; Nishiyama, Y.; Iwata, T.; Wada, M. X-ray crystal structure of anhydrous chitosan at atomic resolution. Biopolymers 2016, 105, 361–368.

(15) Macrae, C. F.; Sovago, I.; Cottrell, S. J.; Galek, P. T. A.; McCabe, P.; Pidcock, E.; Platings, M.; Shields, G. P.; Stevens, J. S.; Towler, M.; Wood, P. A. Mercury 4.0: From Visualization to Analysis, Design and Prediction. J. Appl. Cryst. 2020, 53, 226–235.

(16) Case, D. A.; Ben-Shalom, I. Y.; Brozell, S. R.; Cerutti, D. S.; Cheatham, T., III; Cruzeiro, V. W. D.; Darden, T. A.; Duke, R. E.; Ghoreishi, D.; Gilson, M. K.; Gohlke, H.; Goetz, A. W.; Greene, D.; Harris, R.; Homeyer, N.; Huang, Y.; Izadi, S.; Kovalenko, A.; Kurtzman, T.; Lee, T. S.; LeGrand, S.; Li, P.; Lin, C.; Liu, J.; Luchko, T.; Luo, R.; Mermelstein, D. J.; Merz, K. M.; Miao, Y.; Monard, G.; Nguyen, C.; Nguyen, H.; Omelyan, I.; Onufriev, A.; Pan, F.; Qi, R.; Roe, D. R.; Roitberg, A.; Sagui, C.; Schott-Verdugo, S.; Shen, J.; Simmerling, C. L.; Smith, J.; Salomon-Ferrer, R.; Swails, J.; Walker, R. C.; Wang, J.; Wei, H.; Wolf, R. M.; Wu, X.; Xiao, L.; York, D. M.; Kollman, P. A. AMBER. 2024.

(17) Romany, A.; Payne, G. F.; Shen, J. Effect of Acetylation on the Nanofibril Formation of Chitosan from All-Atom De Novo Self-Assembly Simulations. Molecules 2024, 29, 561.

(18) Guvench, O.; Greene, S. N.; Kamath, G.; Brady, J. W.; Venable, R. M.; Pastor, R. W.; Mackerell, A. D. Additive Empirical Force Field for Hexopyranose Monosaccharides. J. Comput. Chem. 2008, 29, 2543–2564.

(19) Guvench, O.; Mallajosyula, S. S.; Raman, E. P.; Hatcher, E.; Vanommeslaeghe, K.; Foster, T. J.; Jamison, F. W.; MacKerell, A. D. CHARMM Additive All-Atom Force Field for Carbohydrate Derivatives and Its Utility in Polysaccharide and Carbohydrate–Protein Modeling. J. Chem. Theory Comput. 2011, 7, 3162–3180.

(20) MacKerell Jr., A.D.; Bashford, D.; Bellott, M.; Dunbrack, R. L.; Evanseck, J. D.; Field, M. J.; Fischer, S.; Gao, J.; Guo, H.; Ha, S.; Joseph-McCarthy, D.; Kuchnir, L.; Kuczera, K.; Lau, F. T. K.; Mattos, C.; Michnick, S.; Ngo, T.; Nguyen, D. T.; Prodhom, B.; Reiher, W. E.; Roux, B.; Schlenkrich, M.; Smith, J. C.; Stote, R.; Straub, J.; Watanabe, M.; Wiórkiewicz-Kuczera, J.; Yin, D.; Karplus, M. All-Atom Empirical Potential for Molecular Modeling and Dynamics Studies of Proteins. J. Phys. Chem. B 1998, 102, 3586–3616.

(21) Jorgensen, W. L.; Chandrasekhar, J.; Madura, J. D. Comparison of simple potential functions for simulating liquid water. J. Chem. Phys. 1983, 79, 926.

(22) Shirts, M. R.; Klein, C.; Swails, J. M.; Yin, J.; Gilson, M. K.; Mobley, D. L.; Case, D. A.; Zhong, E. D. Lessons learned from comparing molecular dynamics engines on the SAMPL5 dataset. J. Comput.-Aid. Mol. Des. 2017, 31, 147–161.

(23) Grønbech-Jensen, N.; Farago, O. A simple and effective Verlet-type algorithm for simulating Langevin dynamics. Mol. Phys. 2013, 111, 983–991.

(24) Chow, K.-H.; Ferguson, D. M. Isothermal-isobaric molecular dynamics simulations with Monte Carlo volume sampling. Comput. Phys. Commun. 1995, 91, 283–289.

(25) Darden, T.; York, D.; Pedersen, L. Particle Mesh Ewald: An N ·log(N) Method for Ewald Sums in Large Systems. J. Chem. Phys. 1993, 98, 10089–10092.

(26) Miyamoto, S.; Kollman, P. A. Settle: An analytical version of the SHAKE and RATTLE algorithm for rigid water models. J. Comput. Chem. 1992, 13, 952– 962.

(27) Roe, D. R.; Cheatham, T. E. PTRAJ and CPPTRAJ: Software for Processing and Analysis of Molecular Dynamics Trajectory Data. J. Chem. Theor. Comput. 2013, 9, 3084–3095.

(28) Humphrey, W.; Dalke, A.; Schulten, K. VMD: Visual molecular dynamics. J. Mol. Graph. 1996, 14, 33–38.

(29) Virtanen, P.; Gommers, R.; Oliphant, T. E.; Haberland, M.; Reddy, T.; Cournapeau, D.; Burovski, E.; Peterson, P.; Weckesser, W.; Bright, J.; van der Walt, S. J.; Brett, M.; Wilson, J.; Millman, K. J.; Mayorov, N.; Nelson, A. R. J.; Jones, E.; Kern, R.; Larson, E.; Carey, C. J.; Polat, İ.; Feng, Y.; Moore, E. W.; VanderPlas, J.; Laxalde, D.; Perktold, J.; Cimrman, R.; Henriksen, I.; Quintero, E. A.; Harris, C. R.; Archibald, A. M.; Ribeiro, A. H.; Pedregosa, F.; van Mulbregt, P.; SciPy 1.0 Contributors SciPy 1.0: Fundamental Algorithms for Scientific Computing in Python. Nat. Methods 2020, 17, 261–272.

(30) Yui, T.; Imada, K.; Okuyama, K.; Obata, Y.; Suzuki, K.; Ogawa, K. Molecular and Crystal Structure of the Anhydrous Form of Chitosan. Macromolecules 1994, 27, 7601–7605.

(31) Stroud, R. M.; Miercke, L. J.; O’Connell, J.; Khademi, S.; Lee, J. K.; Remis, J.; Harries, W.; Robles, Y.; Akhavan, D. Glycerol Facilitator GlpF and the Associated Aquaporin Family of Channels. Curr. Opin. Struct. Biol. 2003, 13, 424–431.

(32) Milo, R.; Phillips, R. Cell Biology by the Numbers, 1st ed.; Garland Science: New York, 2015.

(33) Tsai, C.-C.; Morrow, B. H.; Chen, W.; Payne, G. F.; Shen, J. Toward Understanding the Environmental Control of Hydrogel Film Properties: How Salt Modulates the Flexibility of Chitosan Chains. Macromolecules 2017, 50, 5946–5952.

(34) Virtanen, P.; Gommers, R.; Oliphant, T. E.; Haberland, M.; Reddy, T.; Cournapeau, D.; Burovski, E.; Peterson, P.; Weckesser, W.; Bright, J.; van der Walt, S. J.; Brett, M.; Wilson, J.; Millman, K. J.; Mayorov, N.; Nelson, A. R. J.; Jones, E.; Kern, R.; Larson, E.; Carey, C. J.; Polat, İ.; Feng, Y.; Moore, E. W.; VanderPlas, J.; Laxalde, D.; Perktold, J.; Cimrman, R.; Henriksen, I.; Quintero, E. A.; Harris, C. R.; Archibald, A. M.; Ribeiro, A. H.; Pedregosa, F.; van Mulbregt, P. SciPy 1.0: Fundamental Algorithms for Scientific Computing in Python. Nat. Methods 2020, 17, 261–272.

(35) Barber, C. B.; Dobkin, D. P.; Huhdanpaa, H. The quickhull algorithm for convex hulls. ACM Trans. Math. Softw. 1996, 22, 469–483.

(36) Okuyama, K.; Noguchi, K.; Miyazawa, T.; Yui, T.; Ogawa, K. Molecular and Crystal Structure of Hydrated Chitosan. Macromolecules 1997, 30, 5849–5855.

(37) Hummer, G.; Garde, S. Cavity Expulsion and Weak Dewetting of Hydrophobic Solutes in Water. Phys. Rev. Lett. 1998, 80, 4193–4196.

(38) Lum, K.; Chandler, D.; Weeks, J. D. Hydrophobicity at Small and Large Length Scales. J. Phys. Chem. B 1999, 103, 4570–4577.

(39) Huang, D. M.; Chandler, D. Temperature and Length Scale Dependence of Hydrophobic Effects and Their Possible Implications for Protein Folding. Proc. Natl. Acad. Sci. U.S.A. 2000, 97, 8324–8327.

(40) Ten Wolde, P. R.; Chandler, D. Drying-Induced Hydrophobic Polymer Collapse. Proc. Natl. Acad. Sci. U.S.A. 2002, 99, 6539–6543.

(41) Liu, P.; Huang, X.; Zhou, R.; Berne, B. J. Observation of a Dewetting Transition in the Collapse of the Melittin Tetramer. Nature 2005, 437, 159–162.

(42) Krone, M. G.; Hua, L.; Soto, P.; Zhou, R.; Berne, B. J.; Shea, J.-E. Role of Water in Mediating the Assembly of Alzheimer Amyloid-β Aβ16-22 Protofilaments. J. Am. Chem. Soc. 2008, 130, 11066– 11072.

(43) Berne, B. J.; Weeks, J. D.; Zhou, R. Dewetting and Hydrophobic Interaction in Physical and Biological Systems. Annu. Rev. Phys. Chem. 2009, 60, 85– 103.

(44) Stillinger, F. H. Structure in Aqueous Solutions of Nonpolar Solutes from the Standpoint of Scaled-Particle Theory. J. Solution Chem. 1973, 2, 141–158.

(45) Romany, A.; Payne, G. F.; Shen, J. Mechanism of the Temperature-Dependent Self-Assembly and Polymorphism of Chitin. Chem. Mater. 2023, 35, 6472–6481.

(46) Shea, J.-E.; Onuchic, J. N.; Brooks, C. L. Probing the Folding Free Energy Landscape of the Src-SH3 Protein Domain. Proc. Natl. Acad. Sci. U.S.A. 2002, 99, 16064–16068.

(47) Tsai, C.-C.; Payne, G. F.; Shen, J. Exploring pH-Responsive, Switchable Crosslinking Mechanisms for Programming Reconfigurable Hydrogels Based on Aminopolysaccharides. Chem. Mater. 2018, 30, 8597–8605.

(48) Smirnov, S. N.; Vlassiouk, I. V.; Lavrik, N. V. Voltage-Gated Hydrophobic Nanopores. ACS Nano 2011, 5, 7453–7461.

(49) Powell, M. R.; Cleary, L.; Davenport, M.; Shea, K. J.; Siwy, Z. S. Electric-Field-Induced Wetting and Dewetting in Single Hydrophobic Nanopores. Nat. Nanotech. 2011, 6, 798–802.

