## Supporting Information for "How Electric Field Remodels the Nanofibril Structure of Chitosan Hydrogels: the Role of Dewetting During Electro-Assembly"

### List of Tables

### List of Figures

S1 Time series of interchain hydrogen bonds and solvent exposure of the chitosan nanofibril with an e-field . . . . . . . . . . . . . . . . . . . . . . . . . . . S-3

S2 Time series of interchain hydrogen bonds and solvent exposure of the chitosan nanofibril without an e-field . . . . . . . . . . . . . . . . . . . . . . . . . S-4

S3 Time series of the intrachain hydrogen bonds with and without an e-fied . . S-5

S4 Time series of the change in the nanofibril volume with and without an e-fieldS-6

S5 Time series of sheet stacking distance (thickness) with and without e-field . S-7

S6 Distribution of sheet growth distance with and without e-field . . . . . . . . . S-8

#### Supplemental figures

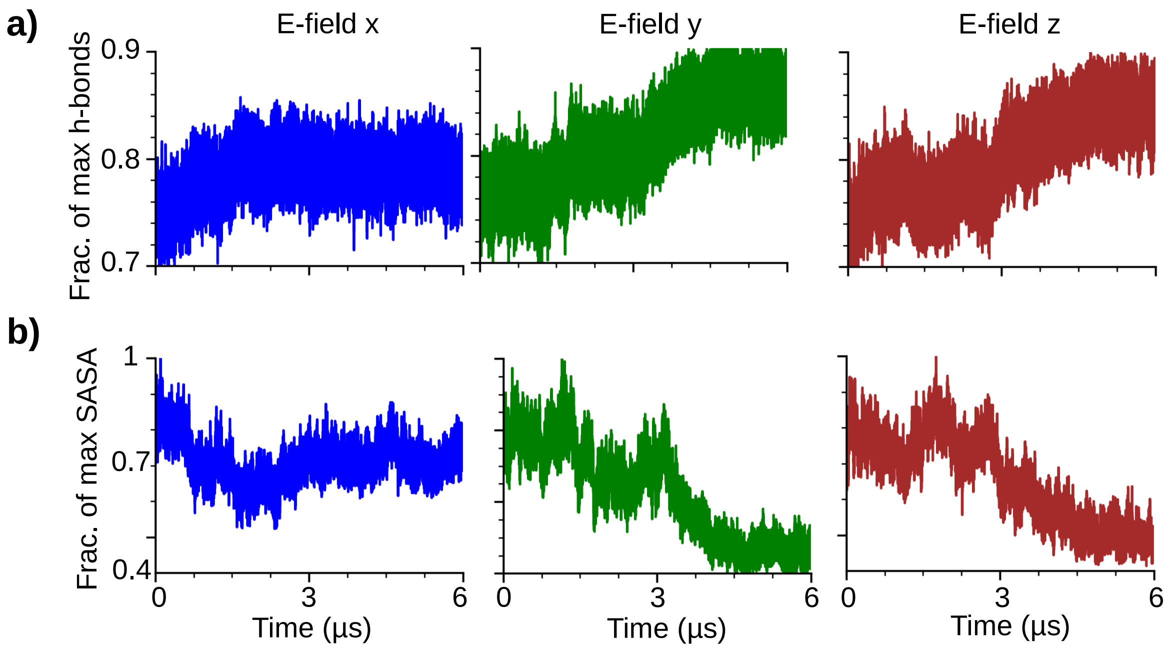


Figure S1: Time series of **(a)** the fraction of max interchain hydrogen bonds and **(b)** solvent accessible surface area (SASA) of the chitosan nanofibril under the electric field in $x$ (left), $y$ (middle), and $z$ (right) direction. A hydrogen bond is considered present if the heavy-atom donor-acceptor distance is below 3.5 Å and the donor-hydrogen-acceptor angle greater than 135°. Note, all 24 chains were used. Fraction is calculated relative to the fibril after energy minimization.

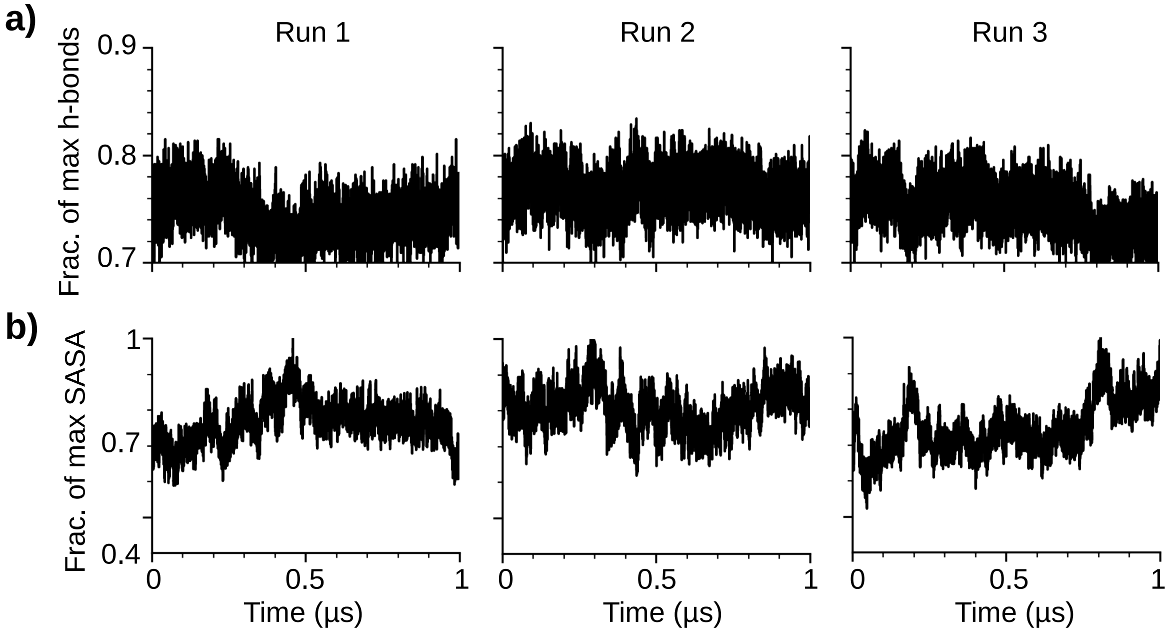


Figure S2: Time series of **(a)** the fraction of max interchain hydrogen bonds and **(b)** the fraction of max SASA in three independent simulations without e-field applied.

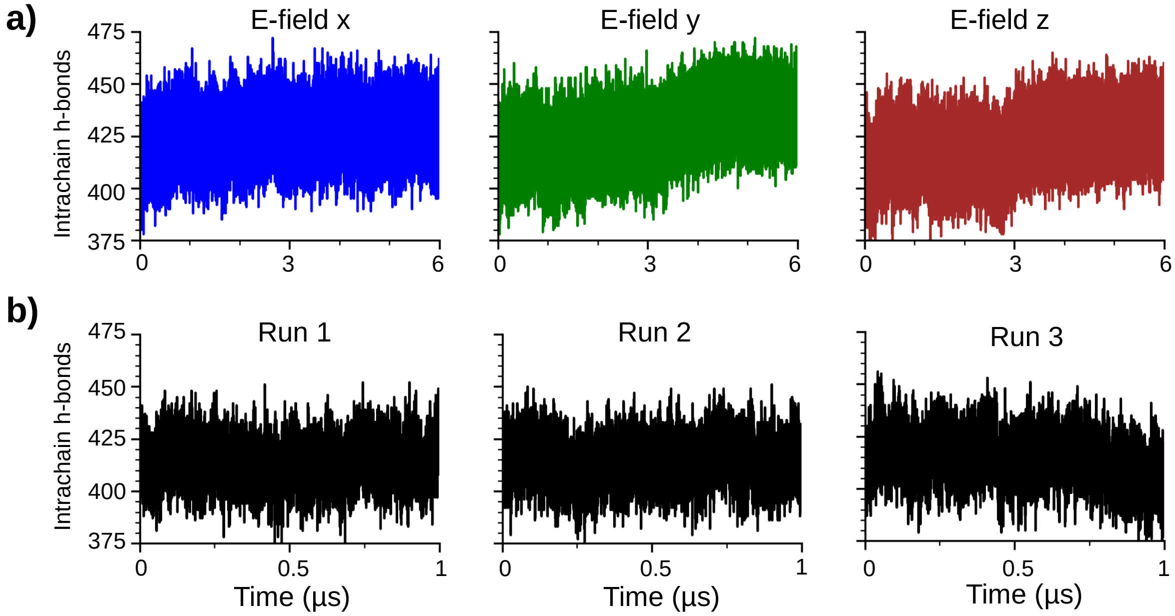


Figure S3: Time series of the total number of intrachain hydrogen bonds in the chitosan nanofibril **(a)** in the simulations with the e-field and **(b)** without an e-field.

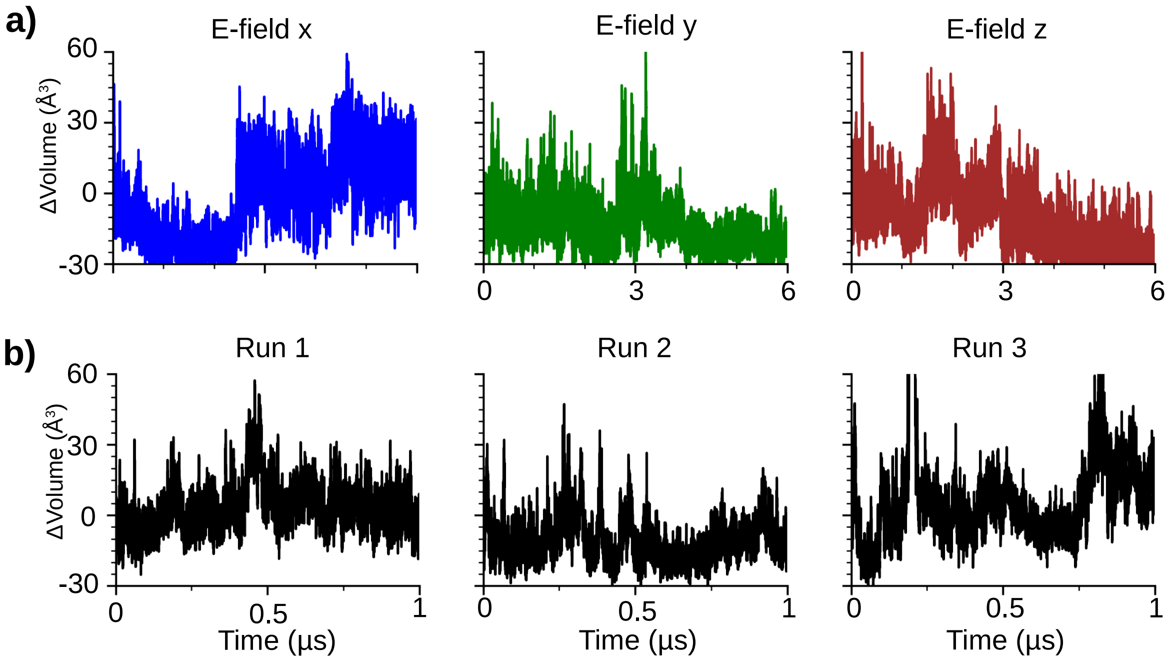


Figure S4: Time series of the change in the nanofibril volume **(a)** with and **(b)** without an e-field applied. The change is calculated relative to the volume of the chitosan nanofibril at the beginning of the production run.

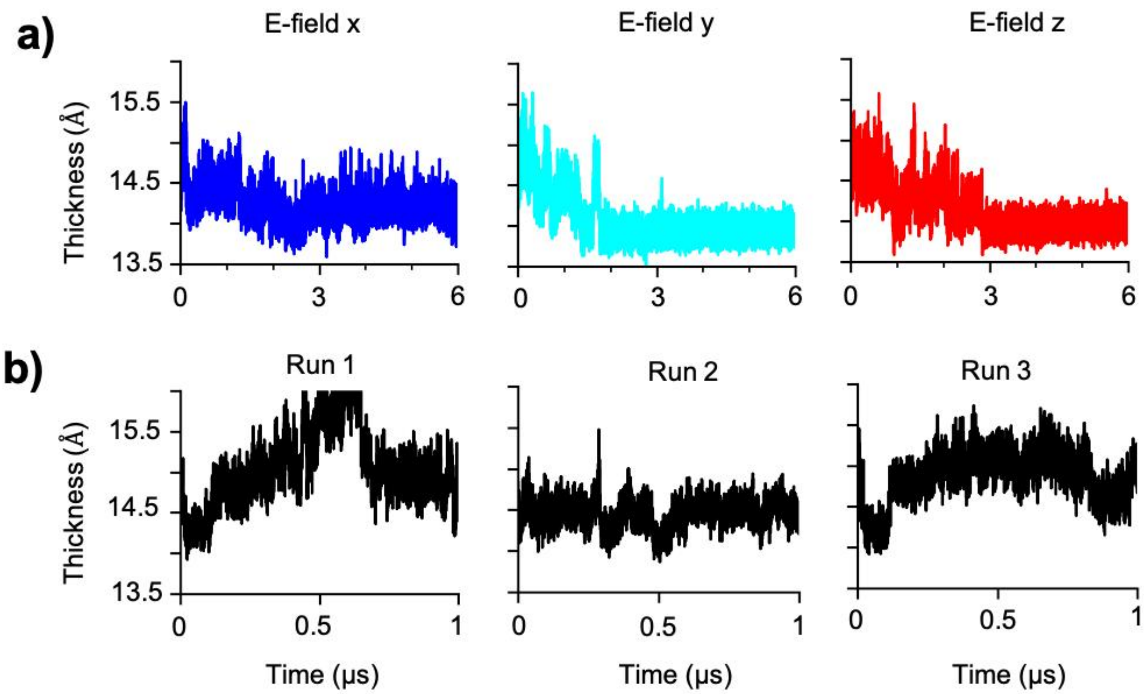


Figure S5: Time series of the change in the nanofibril Thickness **(a)** with and **(b)** without an e-field applied. The thickness is calculated as the distance between center chains in sheet 1 and sheet 4.

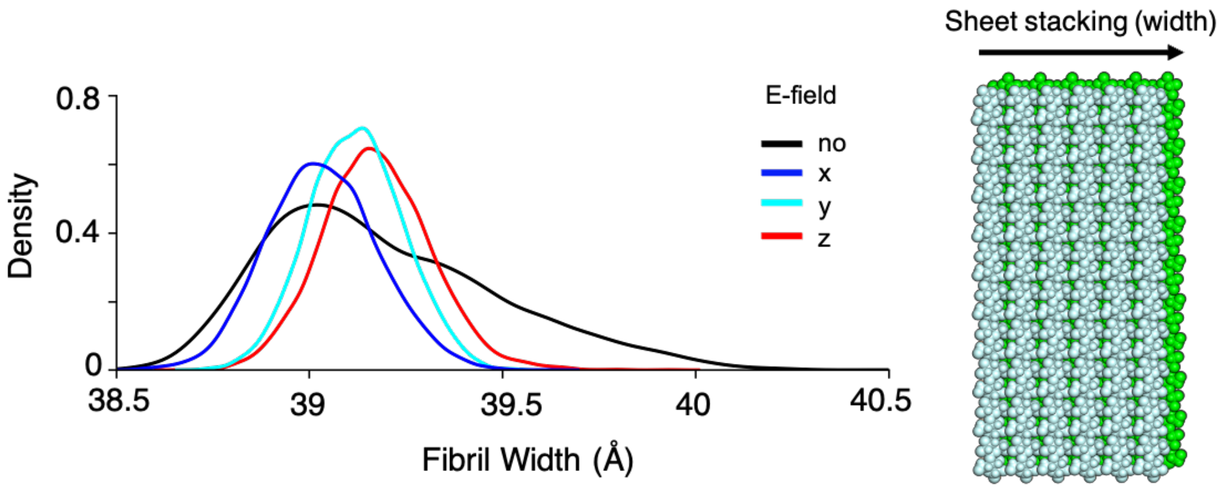


Figure S6: Probability distribution (left) of the distance along the sheet growth direction with and without an e-field applied. It is calculated as the center of mass distances between the two end chains of the inner sheets. Chitosan nanofibril (right) showing the distance illustrating the distance measured.
